# Retrovirus-Like Gag Protein Arc/Arg3.1 is Involved in Extracellular-Vesicle-Mediated mRNA Transfer between Glioma Cells

**DOI:** 10.1101/2023.04.11.536339

**Authors:** Aya Al Othman, Dmitry Bagrov, Julian M Rozenberg, Olga Glazova, Gleb Skryabin, Elena Tchevkina, Alexandre Mezentsev, Mikhail Durymanov

## Abstract

Activity-regulated cytoskeleton-associated (Arc) protein is expressed in neural tissue of vertebrates, where it plays a pivotal role in modulation of synaptic communication. In addition, Arc protein forms capsid-like particles, which can encapsulate and transfer mRNA in extracellular vesicles (EVs) between neurons, that could modulate synaptic function and plasticity. Glioma cell networks actively interact with neurons *via* paracrine signaling and formation of neurogliomal glutamatergic synapses that contribute to cancer cell survival, proliferation, and invasion. Here, we revealed that Arc is expressed in human glioma cell lines, which can produce EVs containing Arc protein and *Arc* mRNA (or “Arc EVs”). Recombinant Arc protein binds to *Arc* mRNA with 1.5-fold higher affinity as compared with control *mCherry* mRNA. Arc EVs from U87 glioma cells internalize and deliver *Arc* mRNA to recipient U87 cells, where it is translated into a protein. Arc overexpression significantly increases EV production, alters EV morphology, and enhances intercellular transfer of highly expressed mRNA in glioma cell culture. These findings indicate involvement of Arc EVs into mRNA transfer between glioma cells that could contribute to tumor progression and affect synaptic plasticity in cancer patients.

## 1. Introduction

Activity-regulated cytoskeleton-associated (Arc) protein, also known as Arg3.1, is a master regulator of synaptic plasticity in mammals. Functionally, Arc protein mediates AMPA receptor internalization in synapses,^1^ thereby playing a significant role in several types of synaptic plasticity, including synaptic scaling, long-term potentiation, and long-term depression.^2–4^ Structural analysis of Arc indicated its remarkable similarity to capsid domain of HIV Gag polyprotein.^5^ It means that mammalian *Arc* gene was domesticated from retroviruses and repurposed during the evolutionary process to participate in intercellular communication in the nervous system. Similar to viral Gag proteins, Arc can form capsid-like particles with its own mRNA. It was determined that Arc capsids transfer mRNA to neurons in extracellular vesicles (EVs) that could be a form of intercellular signaling to control neuroplasticity.^6^

At the same time, EV exchange is an important hallmark of cancer. Intercellular EV transfer accelerates epithelial-to-mesenchymal transition and invasion processes,^7^ partly mediates immunosuppression in different tumors,^8,9^ promotes drug resistance development,^10^ and significantly contributes to a pre-metastatic niche formation.^11,12^ EVs participate in cell-to-cell delivery of numerous biomolecules including different membrane-associated receptors, cell adhesion molecules, transcriptional factors and other proteins, miRNA, mRNA, lncRNA, and DNA.^13^ Involvement of EV exchange into progression of glioma tumors has been reported in multiple studies.^14,15^

In the light of the recent findings in the role of Arc in EV exchange between neurons, we predicted that this protein could be involved in a similar process of EV transfer in glioma tumors. Here, we show expression of Arc protein in several human glioma cell lines. Our results demonstrate the presence of Arc protein in a complex with its own mRNA in EVs from U87 glioma cells. Isolated Arc EVs can deliver *Arc* mRNA to new U87 cells, resulting in its further translation. In addition, Arc overexpression enhances mRNA intercellular transfer in glioma cell culture as compared with Arc KO cells. Thus, our study suggests a new mechanism of mRNA transfer in glioma tumors that could potentially contribute to cancer cell invasion, survival, and development of drug resistance. Moreover, observed phenomenon of dysregulated *Arc* expression in glioma cells might be helpful for understanding of cognition impairment in glioma patients.

## 2. Materials and methods

### 2.1. Plasmids

pCMV6-AC-GFP was purchased from (Origene, RG204129). pHAGE-CMV-eGFP (Addgene #196046), lentiCRISPRv2 (Addgene plasmid #52961), and Addgene’s packaging plasmids psPAX2 (Addgene #12260), pMD.2G (Addgene #12259), pCMV-dR8.91 (Life Science Market) and pHAGE-CMV-RFP (Addgene) were kind gifts from Drs. Olga Glazova and Julian Rozenberg, Moscow Institute of Physics and Technology.

### 2.2. Cell culture

HEK293FT (CRL-3216) (hereafter, “HEK293”), U87-MG (HTB-14), and LN229 (ATCC, CRL-2611) cells were obtained from the American Type Culture Collection (ATCC). U251-MG (09063001) cells were obtained from the European Collection of Authenticated Cell Cultures (ECACC). All cell lines were cultured in Dulbecco’s modified Eagle’s medium (DMEM) (Gibco, Grand Island, NY) supplemented with 10 % fetal bovine serum (FBS) (Gibco, Grand Island, NY) and 1 % penicillin–streptomycin (Invitrogen, Waltham, MA). The cells were incubated at 37L in a humidified 5 % CO_2_ atmosphere. All experiments were performed with mycoplasma-free cells.

### 2.3. EV isolation

Exosomes derived from different cell lines were isolated from 90 mL of serum-free DMEM culture medium containing around 2 × 10^7^ cells after 72-h exposure by ultracentrifugation as described previously.^16^ Briefly, the harvested conditioned medium was subsequently centrifuged at 300 × g for 10 min, 2,000 × g for 10 min, and 10,000 × g for 30 min to remove cells and large debris. Supernatant was collected and filtered through 0.45 µm filters (Merck Millipore) to remove contaminating apoptotic bodies and cell debris. Then, the supernatant was ultracentrifuged at 100,000 × g at 4°C for 90 min using Beckman Coulter Optima™ XE-90 Ultracentrifuge (Beckman Colter, Brea, CA) equipped with Type 45 Ti rotor. The supernatant was gently removed, and crude EV-containing pellets were resuspended in 5 mL of ice-cold PBS and pooled. A second round of ultracentrifugation, 100,000 × g at 4°C for 90 min, was carried out using Type 70 Ti rotor. The resulting EV pellet was resuspended in 30 µL of PBS (stock suspension) and stored at −80°C until use.

### 2.4. LV production and transduction

The Arc-GFP sequence was cloned into the lentivector “plenti-Arc-GFP” through Gibson assembly (for further details, see Supplementary Methods). Arc-GFP, GFP and RFP lentiviruses were produced in HEK293 cells co-transfected with pMD.2G, pCMV-dR8.91, and lentiviral transfer plasmids plenti-Arc-GFP, pHAGE-CMV-eGFP and pHAGE-CMV-RFP, respectively. Co-transfection was carried out using 1 mg mL^-1^ of transfection reagent polyethyenimine (PEI) (Sigma-Aldrich, St. Louis, MO). The supernatant of the transfected cells was collected 48 h later and filtered through a 0.45 μm pore-size filter (Merck Millipore). For viral infection, different human glioma cell lines and HEK293 cells were seeded at 50 % confluency in 6-well plates. On the next day, the virus-containing supernatants from HEK293 cultures mixed with polybrene (Sigma, St. Louis, MO) at a final concentration of 8 mg mL^-1^ were added to the cells in each well. LV-containing supernatants were replaced with a fresh medium after 24 h, followed by 48-h cultivation. Further cell sorting using BioRad S3e Cell Sorter (Biorad, Hercules, CA) resulted in stable Arc-GFP-expressing cells, hereafter referred to as Arc-GFP HEK293, Arc-GFP U87, Arc-GFP U251, and Arc-GFP LN229.

### 2.5. Transmission electron microscopy

Transmission electron microscopy (TEM) was used to visualize isolated EVs. The formvar-carbon grids (Ted Pella, USA) were treated using a glow discharge device Emitech K100X (Quorum Technologies, UK) to increase the sample adsorption onto the carbon surface. The treatment time was 45 s, the current was 25 mA. The samples were applied onto the grids and incubated for 2 min. Next, the grids were blotted, stained with 1 % uranyl acetate solution, blotted again and dried. The images were obtained using a JEM-1400 electron microscope (Jeol, Japan) operating at 120 kV. Obtained images were analyzed in terms of mean EV diameter and particle density per µm^2^.

### 2.6. Nanoparticle tracking analysis (NTA) of the isolated EVs

The size distribution of EVs was determined by NTA using NanoSight LM10 HS instrument equipped with NanoSight LM14 unit (Malvern Panalytical Ltd., Malvern, UK), LM 14C (405 nm, 65 mW) laser unit, and a high-sensitivity camera with a scientific CMOS sensor (C11440-50B, Hamamatsu Photonics, Hamamatsu City, Japan). All measurements were performed in accordance with ASTM E2834–12(2018) using the camera and video processing parameters optimized for EV measurement. Each sample was diluted with particle-free PBS down to a concentration of about 1.5 × 10^8^ particles mL^-1^. Twelve videos 60 s long each were recorded and processed using NTA software 2.3 build 33 (Malvern Panalytical Ltd.). The results from all measurements were combined to obtain a particle size histogram and the total particle concentration corrected for the dilution factor using the NTA software feature.

### 2.7. Western blotting

Cell lysates or EV stock suspensions containing equal amounts of total protein (15 μg) were mixed with loading buffer, boiled for 5 min, separated by denaturing 12.5 % SDS-polyacrylamide gel and transferred to Amersham™ Hybond™ 0.45 μm PVDF membrane (GE Healthcare, UK). The membrane was blocked for 1 h with a blocking buffer (Bio-Rad) under gentle agitation and incubated overnight with antibodies against Arc/Arg3.1 (TA349500, OriGene), CD9 (ab236630, Abcam), CD63 (ab271286, Abcam), CD81 (ab59477, Abcam) or GAPDH (ab8245, Abcam) as a reference. Then, the membrane was washed twice with TBS-T and incubated with HRP conjugated secondary antibody (ab205718, Abcam) at room temperature for 1 h, followed by several washings with TBS-T and deionized water. Protein bands were visualized by ChemiDoc XRS+ imaging system (Bio-Rad, Hercules, CA) using chemiluminescence mode. The signal intensities were normalized to one of GAPDH. The quantification was performed using Quantity One software (Bio-Rad, Hercules, CA, USA).

### 2.8. BLItz analysis of Arc protein binding affinity to mRNA

Binding affinity of Arc protein to mRNA was measured using a bio-layer interferometer BLItz Pro (ForteBio, Pall Life Sciences, NY). Biotinylated mRNAs (Supplementary Methods), *Arc* mRNA and *mCherry* mRNA, were dissolved in sterile-filtered assay PBS buffer (pH 7.4) containing 0.5 mg mL^-1^ of bovine serum albumin (BSA) and 0.002 % Tween 20 to reduce non-specific interactions. The assays were carried out at 25°C in a drop holder and a sample volume of 5 μL. Streptavidin-coated biosensors were pre-equilibrated and hydrated in a 96-well plate for 10 min. As a starting point, the High Precision Streptavidin biosensors were equilibrated in the reaction buffer for 30 s, followed by loading with biotinylated mRNAs at concentration of 50 µg mL^-1^ for 110 s. After the loading step, the biosensors were washed with the assay buffer to establish a new baseline for 45 s. Then, biosensors were exposed to recombinant Arc protein (TP304129, OriGene) solution in the assay buffer at a concentration of 30 µg mL^-1^ of for 130 s. The dissociation of the complexes was monitored further during biosensor washing with the assay buffer. Binding kinetics analysis was performed using ForteBio Data Analysis software.

### 2.9. Inhibitory analysis of EV uptake by U87 cells

U87 cells were grown in 24-well plates at a density of 1 × 10^5^ cells per well for 24 h. Before the addition of fluorescently labeled EVs (the details of labeling are described in Supplementary Methods), the cells were preincubated with different inhibitors of endocytosis for 30 min. For inhibition of clathrin-dependent endocytosis, the cells were treated with 10 μM CPZ. To suppress lipid-raft-dependent endocytosis and macropinocytosis, the cells were treated with 30 μM NYS and 5 μM CytD, respectively. Then, the cells were incubated with PKH26-labeled EVs (1:70 dilution of the stock suspension) in DMEM for 6 h in the presence of the inhibitors. As a positive control, the cells were incubated in DMEM containing PKH26-labeled EVs without any pharmacological treatment. The negative control was the non-treated cells. Unbound particles were removed by washing with PBS. The cells were detached with 0.25 % trypsin and subjected to flow cytometry analysis using CytoFLEX flow cytometer (Beckman Coulter, Brea, CA). Per sample, 10,000 events were gated.

### 2.10. Cell treatment with EVs

Arc-GFP U87 cells were cultured in FBS-free medium for 72 h. The conditioned medium was collected, and EVs were isolated by ultracentrifugation as described above. The U87 recipient cells were seeded on the day before the treatment at confluency of 50 %, and then were cultured for 48 h and 72 h in fresh medium containing EVs from Arc-GFP U87cells (1:70 dilution of the stock suspension). To detect *Arc* and *GFP* expression level, fluorescent imaging and qRT-PCR was conducted.

### 2.11. RNA extraction and qRT-PCR

RNA was purified using ExtractRNA (Evrogen, Moscow, Russia) according to the manufacturer’s protocol. The prepared samples were subjected to spectral analysis. If the absorption ratio A260/A280 was lower than 2.0, the samples were repurified using CleanRNA Standard (Evrogen, Moscow, Russia). cDNA was synthesized by random priming from 1 μg of total RNA using the MMLV RT kit (Evrogen, Moscow, Russia). These samples were subjected to qPCR using CFX96 Real-Time PCR Detection System (Bio-Rad, Hercules, CA, USA) with the primers as follows: human Arc, 5′-CTGAGCCACCTAGAGGAGTACT-3′ and 5′-AACTCCACCCAGTTCTTCACGG-3′, TurboGFP, 5′-CCCGCATCGAGAAGTACGAG-3′ and 5′-GCGGATGATCTTGTCGGTGA-3′ and human GAPDH, 5′-CATGTTCGTCATGGGGTGAACCA-3′ and 5′-AGTGATGGCATGGACTGTGGTCAT-3′.

Amplification of GAPDH was performed for each reverse-transcribed sample as an endogenous quantification standard. The results were analyzed using CFX Manager software supplied by the manufacturer. All samples were run in triplicate.

### 2.12. Co-culture experiment

To detect a contribution of Arc protein to intercellular mRNA transfer, Arc knockout (KO) U87 cell line was obtained (for further details, see Supplementary Methods). Direct co-culture of Arc-GFP U87 with *Arc* KO/RFP U87 cells and *Arc* KO/GFP U87 with *Arc* KO/RFP U87 cells were performed in 6-well plates for 72 h. Initially, 200,000 GFP and RFP-expressing cells in a 6:4 ratio were added per well. The cells were imaged using by AxioVert.A1 microscope (Zeiss, Oberkochen, Germany) equipped with a ×20/0.6 objective lens. Then, all co-cultured cells were harvested and analyzed for GFP and RFP fluorescence using CytoFLEX flow cytometer (Beckman Coulter, Brea, CA). Per sample, 10,000 events were gated. Gene transfer was indicated by the percentage of cells simultaneously expressing GFP and RFP.

### 2.13. Statistical analysis

The statistical data analysis was carried out using Graphpad Prism 5 (GraphPad Software Inc., San Diego, CA) software. The data are presented as mean ± standard deviation (SD). Each experiment was performed as minimum in triplicate. To determine the statistical significance of the differences between two groups, the nonparametric Mann-Whitney U-test was performed. The value p < 0.05 was considered to indicate a statistically significant difference.

## 3. RESULTS

### 3.1. Arc protein is expressed in glioma tumor tissue and in glioma cell lines

It is known that Arc protein is expressed in a limited number of cell types including skin-migratory dendritic cells,^17^ neurons,^4^ and glial cells.^18^ It was also found that Arc protein could be produced by neuroblastoma cancer cells.^19^ We found here that Arc protein is abundantly expressed in several glioma cell lines, including U87, LN229 and U251, as shown by western blot (Figure 1A).

**Fig. 1.**
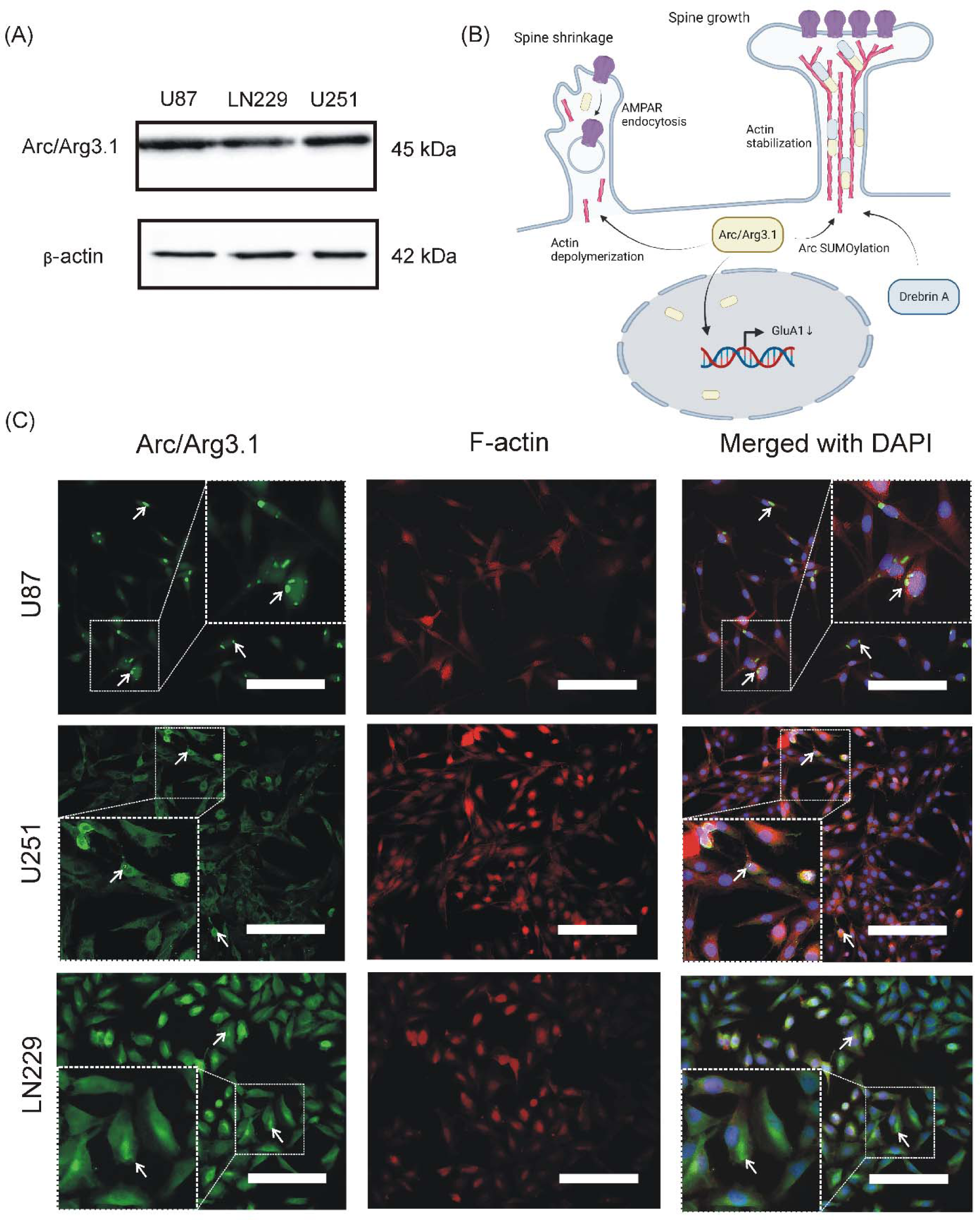
Arc/Arg3.1 expression and subcellular distribution in glioma cells. (A) As shown by immunoblotting, Arc protein is abundantly expressed in glioma cell lines including U87, LN229, and U251. (B) The scheme of Arc protein function and subcellular distribution in neurons. (C) ICC analysis of Arc protein (green) subcellular distribution in glioma cell lines U87, LN229, and U251. Arc was diffusively distributed throughout the cytoplasm volume. Sometimes Arc showed localization in the cell nuclei (blue) and formed protein clusters (white arrows) in cytoplasm nearby actin filament collections (red). The scale bar is 100 µm.

At the subcellular level, Arc protein is non-uniformly distributed in neurons. Upon high frequency stimulation, Arc accumulates in the cell nucleus and regulates PML-dependent GluA1 transcription.^20^ Cytoplasmic Arc *via* interactions with AP-2, endophilin 3, and dynamin 2 mediates endocytosis of AMPARs and dendritic spine shrinkage. SUMOylation of Arc induces its complex formation with drebrin A, resulting in stabilization of actin in the area of the spine and synapse strengthening (Figure 1B).^21^

To elucidate subcellular localization of endogenous Arc protein in U87, LN229, and U251 glioma cell lines, we performed immunocytochemistry (ICC) analysis (for further details, see Supplementary Methods). The cells were stained with fluorescently labeled phalloidin to visualize filamentous actin and with anti-Arc antibodies. It has been shown that in all three cell lines Arc was diffusively distributed throughout the cytoplasm volume. However, sometimes it demonstrated nuclear localization and formed protein clusters in cytoplasm nearby actin filament collections (Figure 1C). Non-uniform subcellular distribution of Arc in glioma cells might indicate involvement of this protein in cytoskeleton organization, vesicular traffic, and nuclear signaling.

### 3.2. Arc protein forms complexes with its own mRNA with increased binding affinity

Besides interaction with multiple protein partners, Arc protein can oligomerize and form capsid-like structures with mRNA. Here, we directly measured binding affinity between Arc protein and mRNA using biolayer interferometry, which is a widely used technique for quantitative analyte detecting as well as binding affinity analysis.^22–24^ In this experiment we also investigated, whether Arc protein discriminates between *Arc* mRNA and other exogenous mRNA (*mCherry*) at the level of binding. For this aim, we immobilized biotinylated *Arc* or *mCherry* mRNA on the streptavidin sensors. On the next step, kinetics of Arc protein binding to the immobilized mRNA and further dissociation was monitored (Figure 2A). Detailed analysis of binding and dissociation curves indicated a more rapid association kinetics for *Arc* mRNA (Figure 2B). Automatic curve-fitting analysis has demonstrated 1.5-fold decreased dissociation constant (K_D_) value for Arc/*Arc* mRNA complex as compared with Arc/*mCherry* mRNA complex indicating the increased specificity of Arc protein to its own mRNA (Figure 2C-E and Table S1).

**Fig. 2.**
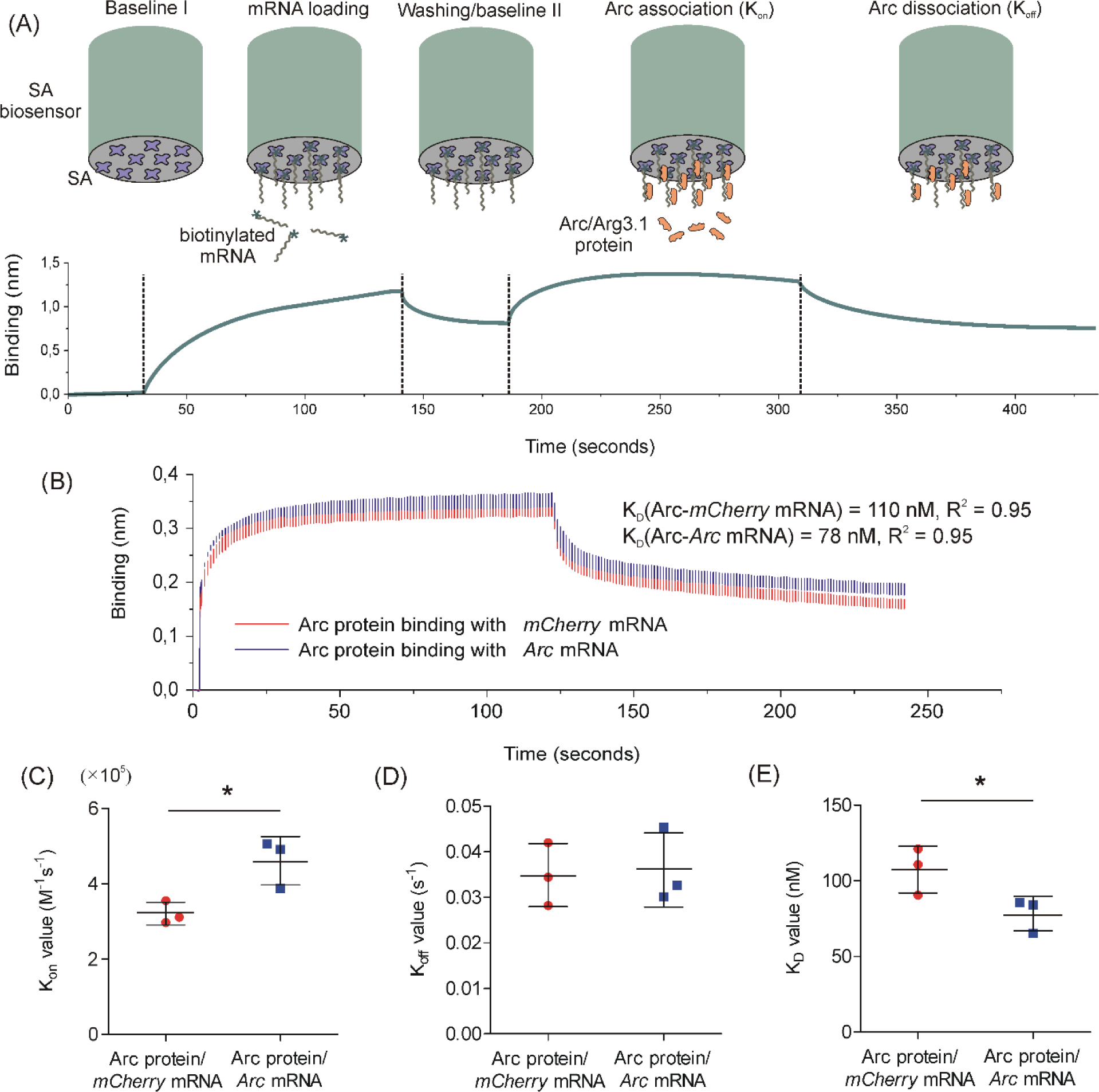
Measurement of Arc/Arg3.1 protein binding affinity to mRNA using biolayer interferometry. (A) The scheme of the measurement procedure, which includes the following steps: 1) incubation of streptavidin (SA) biosensors in PBS to set baseline I; 2) loading of the biosensor with biotinylated mRNA; 3) washing of biotinylated mRNA and the setting baseline II; 4) binding with Arc protein; 5) dissociation of Arc protein. (B) Fitted experimental Arc-mRNA association and dissociation curves shown as means with SD error bars. Calculated values of (C) K_on_, (D) K_off_, and (E) apparent K_D_ for Arc protein/*mCherry* mRNA and Arc protein/*Arc* mRNA shown as means ± SD. * p < 0.05, Mann-Whitney U test.

### 3.3. Arc protein and Arc mRNA are present in U87 produced EVs

Arc/mRNA capsids can be incorporated into EVs and released from neurons. We hypothesized that a similar process may occur in glioma cells. To evaluate the ability of glioma cells to produce EVs containing Arc protein and *Arc* mRNA, we investigated characteristics of EVs isolated from U87 cells. Usually, EV population includes two types of vesicles such as exosomes and microvesicles (MVs). Exosomes with an average size of < 120 nm are formed within multi-vesicular bodies (MVB) and are released from the cells by exocytosis. MVs are larger than exosomes, and they are released from the cells by pinching off the cell surface.^25^

NTA analysis of the U87-derived EVs has shown that their mean diameter was 130 ± 10 nm indicating the presence of both exosome and MV fractions (Figure 3A). Immunoblotting analysis indicated the exosome-specific protein markers such as CD9, CD63, and CD81 (exosome-specific tetraspanins) associated with glioma EVs (Figure 3B). Besides tetraspanins, EVs from glioma cells contained a small amount glycolytic enzyme glyceraldehyde-3-phosphate dehydrogenase (GAPDH), which is directly involved in generation of intraluminal vesicles in multi-vesicular bodies and in exosome clustering.^26^ It was also found that Arc protein is also present in the U87-derived EVs (Figure 3B).

**Fig. 3.**
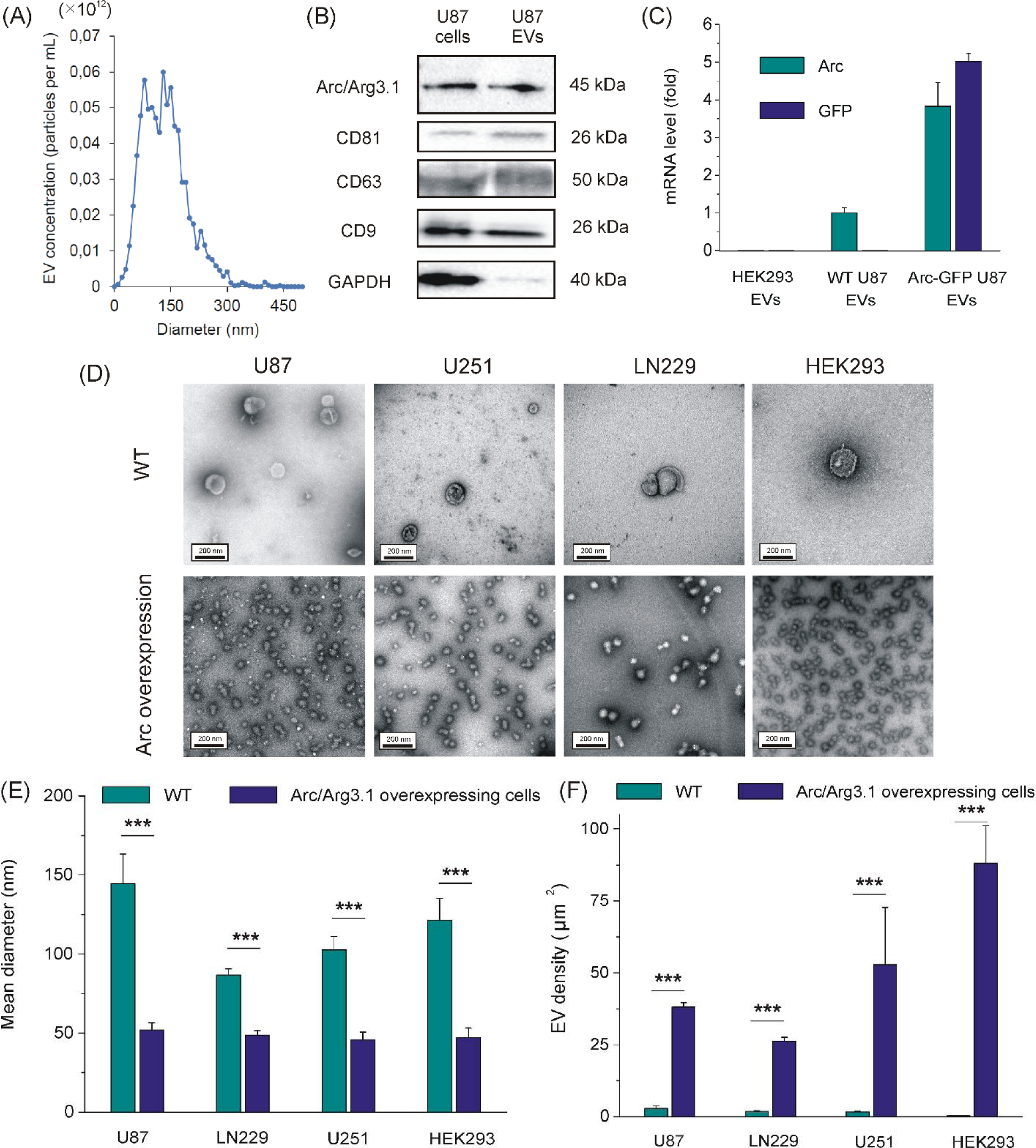
Characterization of Arc EVs. (A) As determined by NTA analysis, Arc EVs demonstrated two-peak size distribution with average diameter of 130 nm. (B) Besides Arc/Arg3.1 protein, immunoblotting analysis indicated the presence of exosome-specific tetraspanins (CD9, CD63, and CD81) and less amount of GAPDH associated with Arc EVs. (C) qRT-PCR analysis of *Arc* and *GFP* mRNA content in EVs obtained from HEK293, WT U87, and Arc-GFP U87 cells. (D) TEM images of EVs, isolated from WT and Arc-overexpressing HEK293, U87, LN229, and U251 cells. The scale bar is 200 nm. Quantitative analysis of TEM images of EVs showed a decrease in the mean diameter of EVs (E) and a significant increase in their production (F) by Arc-overexpressing cells. Data are shown as means ± SD. *** p < 0.001, Mann-Whitney U test.

Once Arc protein binds with *Arc* mRNA, it was reasonable to predict the presence of this mRNA in EVs. qRT-PCR analysis has shown that U87-derived EVs contain *Arc* mRNA, whereas no *Arc* mRNA was found in EVs from HEK293 cells, which do not express Arc protein (Figure 3C). It should be noted that EVs from Arc-GFP overexpressing U87 cells contained 4-fold higher amount of *Arc* mRNA as compared to WT U87. Moreover, they displayed an elevated *GFP* mRNA level indicating the ability of exogenously expressed Arc protein to be incorporated in EVs (Figure 3C).

Once Arc protein forms capsid-like structures, we used Arc-GFP overexpressing glioma cells for evaluation of Arc contribution to the EV production and morphology. We used TEM for the analysis of EVs from Arc WT and Arc-GFP overexpressing glioma cell lines U87, LN229, and U251. As a control, we used EVs from non-expressing Arc WT HEK293 cells and Arc-GFP-transfected HEK293 cells. We found that Arc overexpression significantly increased EV production rate by all glioma cell types and HEK293 cells. Arc overexpression also led to a 2-3-fold decrease of their mean size. Moreover, EVs from Arc-overexpressing cells acquired “virus-like” morphology (Figure 3D-F). Thus, our data suggests the involvement of Arc protein in production of Arc EVs with morphology, different from exosomes and microvesicles.

### 3.4. Arc EVs internalize to U87 cells via macropinocytosis and mediate mRNA translation in recipient U87 cells

We found that glioma cells produce EVs containing Arc protein and *Arc* mRNA (Arc EVs). On the next step, we studied an ability of Arc EVs to deliver mRNA to recipient glioma cells. EVs mediate delivery of multiple molecular cargos to different cells. Uptake kinetics of Arc EVs was determined by flow cytometry. For this purpose, Arc EVs were fluorescently labeled with PKH26 dye and added to U87 cells (Figure 4A). PKH26 is a lipophilic dye, which stains an EV membrane. It was found that Arc EVs gradually internalized by U87 cells over time (Figure 4A,B).

**Fig. 4.**
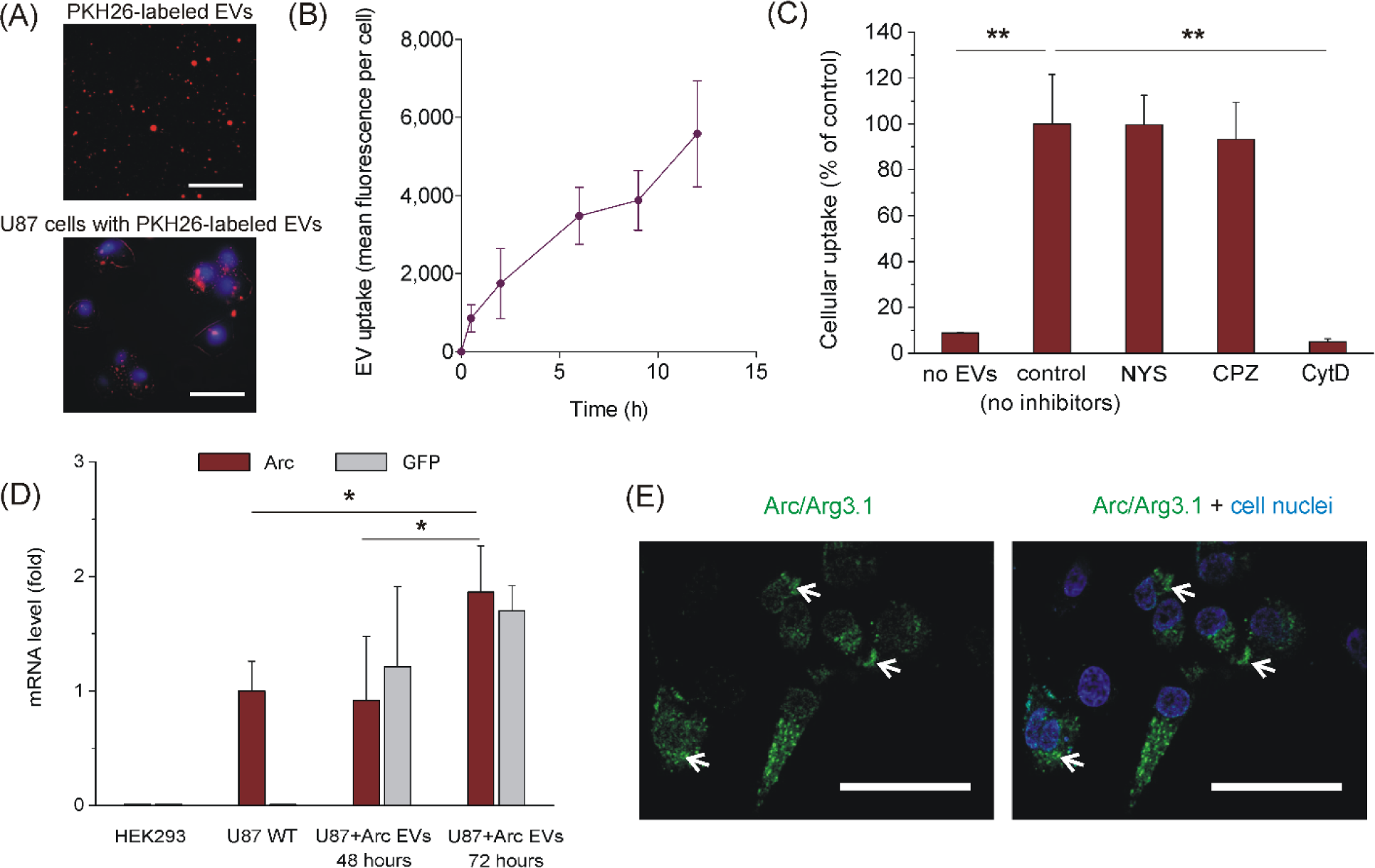
Arc EV uptake and mRNA transfer. (A) Photographs of PKH-26-labeled Arc EVs (red) and Hoechst 33342-stained U87 cells (blue cell nuclei) after 6-h incubation with fluorescent EVs. The scale bars are 10 µm and 20 µm for upper and lower images, respectively. (B) Kinetics of Arc EV uptake by U87 cells shown as mean fluorescence per cell. (C) Inhibitory analysis of Arc EV endocytic routes. (D) qRT-PCR analysis of *Arc* and *GFP* mRNA content in recipient U87 glioma cells treated with Arc EVs from Arc-GFP overexpressing U87 cells. Data are shown as means ± SD. * p < 0.05, ** p < 0.01, Mann-Whitney U test. (E) An image of U87 cells after 72-h incubation with Arc EVs from Arc-GFP overexpressing U87 cells. White arrows show Arc-GFP protein clusters in recipient cells. The scale bar is 30 µm.

It should be noted that cellular uptake can be mediated through multiple routes including phagocytosis/macropinocytosis, clathrin-mediated endocytosis, and lipid-raft dependent uptake. To determine the preferred cellular uptake pathway, we used pharmacological inhibitors cytochalasin D (CytD), chlorpromazine (CPZ), and nystatin (NYS) to evaluate uptake contribution of macropinocytosis, clathrin-mediated, and lipid-raft dependent endocytosis, respectively. For this experiment, we determined maximal non-toxic concentrations of each inhibitor assessed using MTT test (Supplementary Methods and Figure S1). We found that 30 µM CPZ, 70 µM NS, and 5 µM CytD did not significantly affect viability of U87 cells. It has been revealed that only CytD significantly inhibited the cell entry of PKH26-labeled Arc EVs (Figure 4C). It means that cellular uptake of Arc EVs by U87 cells is mediated by macropinocytosis route.

We found that treatment of recipient U87 cells with Arc EVs isolated from Arc-GFP-transfected U87 cells resulted in a significant increase in the level of Arc and GFP expression upon 48 and 72 h of incubation (Figure 4D). We also found the appearance of Arc-GFP expression in recipient U87 cells (Figure 4E), which demonstrated a similar subcellular distribution as an endogenous Arc protein. These data suggest that Arc EVs are involved in intercellular mRNA transfer between glioma cells.

### 3.5. Arc overexpression enhances mRNA intercellular transfer in glioma cell co-culture

Once Arc EVs are involved into intercellular mRNA transfer, we aimed to evaluate their contribution in comparison with Arc-independent routes of mRNA transfer. For this experiment we produced Arc KO U87 cells (Supplementary Methods and Figure 5A,B) with permanent RFP expression (Arc KO/RFP U87). Then, we used these cells for the establishment of two co-culture systems (Figure 5C). In the first case, we co-cultured these cells with Arc KO U87 GFP-expressing cells (Arc KO/GFP U87). We expected that *GFP* and *RFP* mRNA transfer in the co-culture of two Arc KO U87 cell lines relies exclusively on non-Arc EV-mediated exchange and tunneling nanotube (TNT)-mediated transfer. The second co-culture system comprising Arc-GFP U87 cells and Arc KO/RFP U87 cells was used to evaluate a contribution of Arc EVs into transfer of *GFP* mRNA (Figure 5C). After 48 h of incubation, we examined a fraction of the cells with simultaneous GFP and RFP expression in the co-cultured populations. Interestingly, GFP and RFP-positive cells were present even in the co-culture of Arc KO cell lines (around 6 %), thereby suggesting the existence of Arc-independent routes of mRNA transfer (Figure 5D,E). At the same time, the percentage of GFP and RFP co-expressing cells in the co-culture of Arc-GFP U87 and Arc KO/RFP U87 cells was 3-fold higher than in Arc KO co-culture (Figure 5D,E). Fluorescent microscopy of these cells has shown the same subcellular distribution of Arc-GFP (Figure 5F) as in Arc WT U87 cells. This result indicates a significant contribution of Arc EVs to intercellular mRNA transfer between glioma cells.

**Fig. 5.**
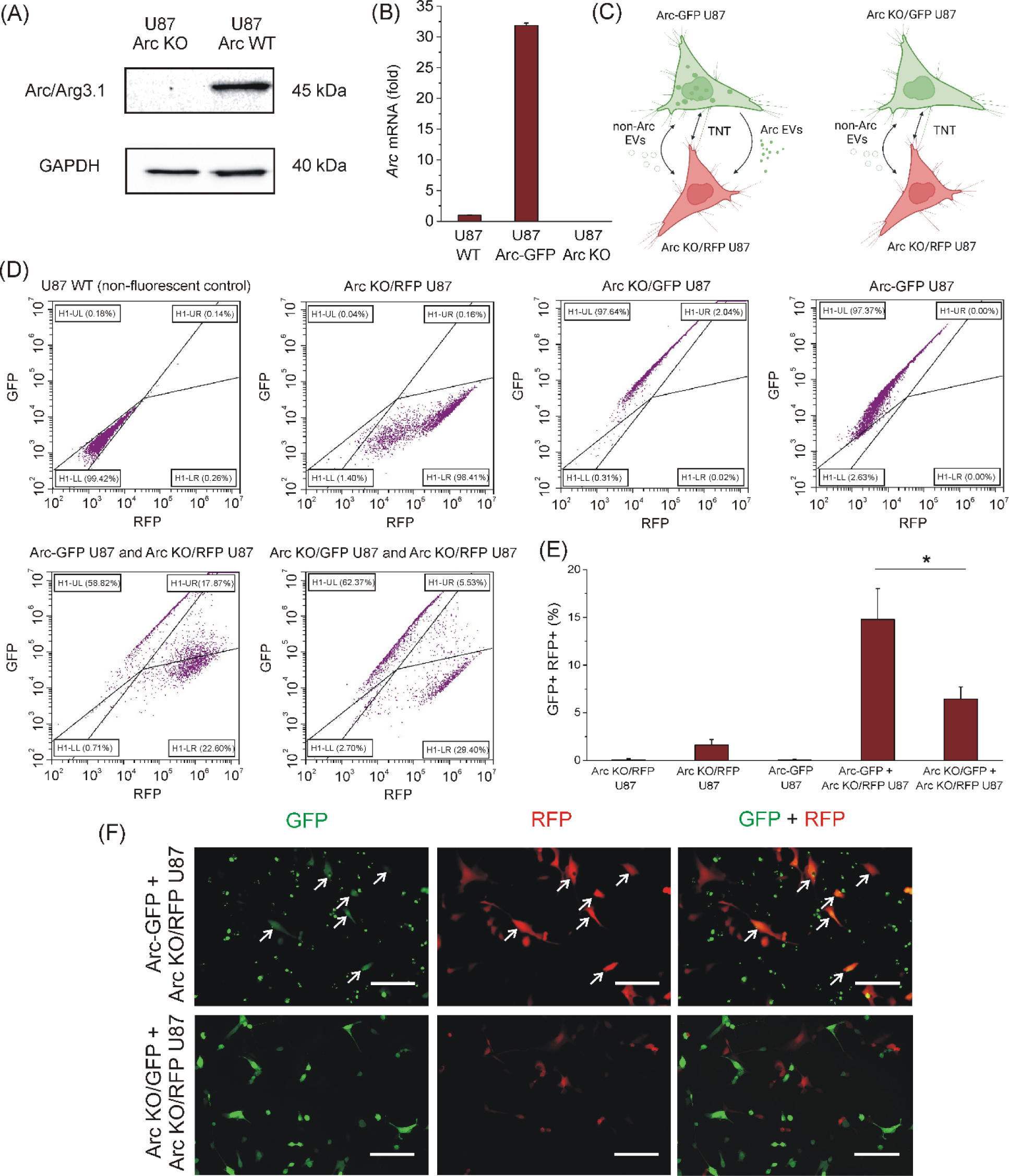
Influence of Arc expression on *GFP* and *RFP* mRNA transfer in glioma cell co-culture. (A) Immunoblotting evaluation of Arc/Arg3.1 protein expression by Arc KO U87 cells. (B) Evaluation of *Arc* mRNA expression level in Arc WT, Arc KO, and Arc-GFP U87 cells. (C) A scheme of mRNA transfer in Arc-GFP + Arc KO/RFP U87 and Arc KO/GFP + Arc KO/RFP U87 cell co-cultures. (D) Flow cytometry analysis of GFP- and RFP-expressing U87 cell lines and their co-cultures. (E) It was found that the percentage of GFP and RFP-positive cells in the co-culture of Arc-GFP U87 and Arc KO/RFP U87 cells was 3-fold higher than in Arc KO co-culture. Data are shown as means ± SD. * p < 0.05, Mann-Whitney U test. (F) Images of Arc-GFP + Arc KO/RFP U87 and Arc KO/GFP + Arc KO/RFP U87 cell co-cultures after 72-h incubation. White arrows indicate GFP and RFP co-expressing cells. The scale bar is 50 µm.

## 4. Discussion

Gliomas are the most common type of brain malignancies.^27^ These tumors are characterized by high clonal and morphological heterogeneity, rapid proliferation, remarkable infiltrative growth, drug resistance, and poor survival rate.^28^ Surprisingly, some of these properties are promoted by interaction of glioma cells with normal neurons. First, neuronal activity promotes the growth of malignant glioma through paracrine secretion of the synaptic protein neuroligin-3, which activates PI3K-Akt-mTOR signaling pathway to induce glioma cell survival and proliferation.^29^ Second, glioma cells directly interact with neurons and form bona fide glutamatergic synapses consisting of a presynaptic neuronal and a postsynaptic gliomal part. Neuronal activity produces currents in postsynaptic glioma cell membrane and intracellular calcium transients in glioma cell network that induces calcium-dependent pro-survival signaling pathways resulting in growth and invasiveness of brain tumors.^30,31^

Once glioma and neural networks can interact *via* synaptic connections, we hypothesized that glioma cells could express *Arc* gene, involved in regulation of synaptic plasticity. We revealed that Arc protein is expressed in several glioma cell lines (Figure 1A). It was found that Arc protein was localized diffusively in cytoplasm of glioma cells or formed protein clusters (Figure 1C) resembling behavior of nascent HIV-1^32^ and M-PMV^33^ Gag polyproteins.

Since Arc forms virus-like particles with mRNA, we first evaluated affinity of Arc protein binding to mRNA. Previously, specificity of Arc protein interaction with different mRNAs was examined using qRT-PCR analysis of the samples obtained from the whole cell bacteria lysate and the Arc protein fraction, co-purified with bound mRNA. It turned out that the ratio between *Arc* mRNA and bacterial *asnA* mRNA in both samples remained the same suggesting that Arc capsids have a little specificity for a particular mRNA.^6^ We used here biolayer interferometry to measure binding affinity of Arc protein binding to *Arc* mRNA versus control *mCherry* mRNA. It was found that pure Arc protein binds to *Arc* mRNA with 1.5-fold higher affinity as compared with *mCherry* mRNA (apparent K_D_ = 78 nM for *Arc* mRNA and 110 nM for *mCherry* mRNA) (Figure 2B-E and Table S1). Interestingly, the value of apparent K_D_ value for Arc-mRNA complexes is of the same order as for HIV-1 Gag polyprotein and viral RNA.^34^

It was found that glioma cells can produce EVs, containing Arc protein and *Arc* mRNA (Figure 3), as was earlier discovered for the neurons.^6^ Although Arc EVs contained such exosome markers as CD81, CD63, CD9, and GAPDH, NTA and SEM analyses displayed a significant size heterogeneity indicating a presence of non-exosome EV fraction in the conditioned medium. However, Arc overexpression markedly increased a small-size EV fraction and total amount of the released EVs (Figure 3D-F). The major EV fraction in Arc-overexpressing glioma cells is around 40-50 nm in diameter that is very similar to earlier reported size of Arc capsids.^6^ At the same time, these vesicles morphologically resemble viral particles rather than exosomes. It means that elevated level of Arc expression enhances EV production and alters vesicle morphology.

Arc EVs can be internalized by neurons *via* clathrin-mediated endocytic pathway.^6^ Our data demonstrated a relatively slow kinetics of EV uptake resulting in saturation at 6 h after EV treatment (Figure 4B) that is not typical for rapid clathrin-mediated endocytosis (CME), which takes from seconds to minutes.^35^ Inhibitory analysis of cellular uptake pathways has shown that fluorescently labeled EVs are internalized exclusively due to micropinocytosis (Figure 4C). Actually, this result might not be contradictory to the data obtained for neurons because the authors used dynasore to inhibit CME.^6^ Dynasore inhibits not only dynamin-dependent processes, including CME, but also suppresses micropinocytosis as well.^36^

RT-PCR analysis has shown that the levels of exogenous Arc and GFP expression significantly increase in recipient U87 cells over time indicating accumulation of exogenous intact *Arc-GFP* mRNA upon Arc EV uptake (Figure 4D). Fluorescent imaging of recipient Arc EV-treated cells demonstrated the same intracellular localization of GFP-tagged Arc protein (Figure 4E) as in Arc WT U87 cells (Figure 1C) suggesting that exogenous *Arc-GFP* mRNA transferred with Arc EVs is translated into a protein. Since Arc protein is involved into EV-mediated mRNA delivery to recipient cells and its overexpression stimulates EV production, we tested whether expression level of Arc in glioma cells affects the rate of mRNA transfer between glioma cells. For this purpose, we compared co-cultures of Arc-GFP-expressing and Arc KO/RFP-expressing U87 cells versus Arc KO GFP-expressing and Arc KO RFP-expressing U87 cells. Flow cytometry analysis of co-cultures indicated an increase in GFP and RFP co-expressing cells in both co-cultures indicating existence of Arc-independent mRNA transfer mechanism (Figure 5C). Possible mechanisms might include Arc-independent EV exchange and mRNA transfer *via* tunneling nanotubes (TNTs), dynamic cytoplasmic bridges between the cells. As was shown earlier, U87 cells could form TNTs.^37^ However, we observed 3-fold higher increase in GFP and RFP co-expressing cells in Arc-GFP-expressing and Arc KO/RFP-expressing U87 co-cultures as compared with Arc KO co-cultures (Figure 5E) that indicates a significant contribution of Arc expression to EV-mediated mRNA transfer.

In conclusion, we revealed that glioma cells can release Arc EVs, which mediate uptake and translation of exogenous mRNA in new glioma cells. This result could have some implications for better understanding intercellular crosstalk in glioma microenvironment.

First, gliomas are genetically heterogenous group of tumors. Once Arc protein is not so highly selective for a particular mRNA, Arc EVs could transfer other abundantly expressed mRNAs between glioma cells. It means that Arc EV-mediated communication between different clones of glioma cells would enrich them with mRNAs encoding oncoproteins that would facilitate development of drug resistance.

Second, glioma-produced Arc EVs could be involved into crosstalk with non-glioma cells in tumor microenvironment. Among them, neurons also produce Arc EVs. It is though that Arc EV transfer between neurons might be involved into control of synaptic function and plasticity^6^ because intracellular Arc, on the one hand, stabilizes expansion of the F-actin network in dendritic spines resulting in morphological enlargement of the synapse,^38^ and on the other hand, Arc is implicated into AMPA receptor uptake and spine elimination.^39^ In this regard, further investigations should determine whether cancer cell-produced Arc EVs interact with neurons and contribute to neurogliomal synaptic interaction, which stimulates calcium entry to glioma cells, activation of calcium-dependent pro-survival signaling pathways, and better resistance to radiotherapy and chemotherapy.^30,31^ Besides neurons, Arc EVs could contribute to mRNA delivery to other non-glioma cells including tumor-associated macrophages and T-cells. Therefore, the contribution of this process to immunosuppression should also be investigated in future.

Next, if EV-mediated mRNA transfer between glioma cells is dependent on Arc, the factors affecting *Arc* expression should also be investigated. *Arc* belongs to immediate early genes that are transcribed in response to synaptic input in active neurons.^4^ Since glioma cell networks also receive synaptic input from neurons, *Arc* expression also could be dependent on neuronal activity. It was also found recently that Arc protein expression in astrocytes can be enhanced in response to lactate.^40^ Glioma tumors often contain hypoxic areas and elevated lactate concentration in tumor tissue.^41^ Therefore, abundant lactate production also might be a factor affecting *Arc* expression in glioma cells.

Finally, different forms of memory including verbal and visuospatial memory are impaired in glioma patients as a consequence of tumor development.^42^ Probably, neurogliomal synapses formation along with dysregulated Arc EV-mediated intercellular signaling in the brain are implicated into cognitive function impairment in glioma patients.

## Supporting information

subblementary

## Acknowledgements

This work was supported by the Russian Science Foundation grant no. 21-74-00019. The experiments were carried out at the User Facilities Center “Electron microscopy in life sciences” at Lomonosov Moscow State University.

## Declaration of competing interest

The authors declare that they have no known competing financial interests or personal relationships that could have appeared to influence the work reported in this paper.

## Author contributions

Mikhail Durymanov and Aya Al Othman developed the concept. Aya Al Othman carried out ICC, western blotting experiments, RT-PCR, cloning, uptake inhibitory analysis, co-culture experiments, and BLItz. Dmitry Bagrov performed TEM experiments. Julian Rozenberg and Olga Glazova contributed to work with cloning, lentiviral transfection, and development of Arc KO U87 cell lines. Gleb Skryabin and Elena Tchevkina contributed to EV isolation and implemented NTA analysis. Mikhail Durymanov, Alexandre Mezentsev, and Aya Al Othman wrote a manuscript. All authors have read and approved the final version of the manuscript.

## Supporting Information

The Supporting Information is available free of charge at …

## Notes

### Competing Interest Statement

The authors have declared no competing interest.

### Summary of Updates

1. the concentration of EVs after isolation from cells (stock) 2. the dilutions of EVs made for each experiment before conducting. 3. authors contribution 4. Acknowledgements

## References

(1) Chowdhury, S.; Shepherd, J. D.; Okuno, H.; Lyford, G.; Petralia, R. S.; Plath, N.; Kuhl, D.; Huganir, R. L.; Worley, P. F. Arc/Arg3. 1 Interacts with the Endocytic Machinery to Regulate AMPA Receptor Trafficking. Neuron 2006, 52 (3), 445–459.

(2) Kedrov, A. V.; Durymanov, M.; Anokhin, K. V. The Arc Gene: Retroviral Heritage in Cognitive Functions. Neuroscience & Biobehavioral Reviews 2019, 99, 275–281. https://doi.org/10.1016/j.neubiorev.2019.02.006.

(3) Fila, M.; Diaz, L.; Szczepanska, J.; Pawlowska, E.; Blasiak, J. MRNA Trafficking in the Nervous System: A Key Mechanism of the Involvement of Activity-Regulated Cytoskeleton-Associated Protein (Arc) in Synaptic Plasticity. Neural Plasticity 2021, 2021, e3468795. https://doi.org/10.1155/2021/3468795.

(4) Zhang, H.; Bramham, C. R. Arc/Arg3.1 Function in Long-Term Synaptic Plasticity: Emerging Mechanisms and Unresolved Issues. Eur J Neurosci 2021, 54 (8), 6696–6712. https://doi.org/10.1111/ejn.14958.

(5) Zhang, W.; Wu, J.; Ward, M. D.; Yang, S.; Chuang, Y.-A.; Xiao, M.; Li, R.; Leahy, D. J.; Worley, P. F. Structural Basis of Arc Binding to Synaptic Proteins: Implications for Cognitive Disease. Neuron 2015, 86 (2), 490–500. https://doi.org/10.1016/j.neuron.2015.03.030.

(6) Pastuzyn, E. D.; Day, C. E.; Kearns, R. B.; Kyrke-Smith, M.; Taibi, A. V.; McCormick, J.; Yoder, N.; Belnap, D. M.; Erlendsson, S.; Morado, D. R. The Neuronal Gene Arc Encodes a Repurposed Retrotransposon Gag Protein That Mediates Intercellular RNA Transfer. Cell 2018, 172 (1–2), 275–288.

(7) Kim, H.; Lee, S.; Shin, E.; Seong, K. M.; Jin, Y. W.; Youn, H.; Youn, B. The Emerging Roles of Exosomes as EMT Regulators in Cancer. Cells 2020, 9 (4), 861.

(8) Xie, Q.-H.; Zheng, J.-Q.; Ding, J.-Y.; Wu, Y.-F.; Liu, L.; Yu, Z.-L.; Chen, G. Exosome-Mediated Immunosuppression in Tumor Microenvironments. Cells 2022, 11 (12), 1946.

(9) Olejarz, W.; Dominiak, A.; Żolnierzak, A.; Kubiak-Tomaszewska, G.; Lorenc, T. Tumor-Derived Exosomes in Immunosuppression and Immunotherapy. Journal of immunology research 2020, 2020.

(10) Bach, D.-H.; Hong, J.-Y.; Park, H. J.; Lee, S. K. The Role of Exosomes and MiRNAs in Drug-Resistance of Cancer Cells. International journal of cancer 2017, 141 (2), 220–230.

(11) Lobb, R. J.; Lima, L. G.; Möller, A. Exosomes: Key Mediators of Metastasis and Pre-Metastatic Niche Formation. In Seminars in cell & developmental biology; Elsevier, 2017; Vol. 67, pp 3–10.

(12) Guo, Y.; Ji, X.; Liu, J.; Fan, D.; Zhou, Q.; Chen, C.; Wang, W.; Wang, G.; Wang, H.; Yuan, W. Effects of Exosomes on Pre-Metastatic Niche Formation in Tumors. Molecular Cancer 2019, 18 (1), 1–11.

(13) Mashouri, L.; Yousefi, H.; Aref, A. R.; Ahadi, A. mohammad; Molaei, F.; Alahari, S. K. Exosomes: Composition, Biogenesis, and Mechanisms in Cancer Metastasis and Drug Resistance. Molecular cancer 2019, 18, 1–14.

(14) Gourlay, J.; Morokoff, A. P.; Luwor, R. B.; Zhu, H.-J.; Kaye, A. H.; Stylli, S. S. The Emergent Role of Exosomes in Glioma. Journal of Clinical Neuroscience 2017, 35, 13–23.

(15) Shi, J.; Zhang, Y.; Yao, B.; Sun, P.; Hao, Y.; Piao, H.; Zhao, X. Role of Exosomes in the Progression, Diagnosis, and Treatment of Gliomas. Medical science monitor: international medical journal of experimental and clinical research 2020, 26, e924023–1.

(16) Doyle, L. M.; Wang, M. Z. Overview of Extracellular Vesicles, Their Origin, Composition, Purpose, and Methods for Exosome Isolation and Analysis. Cells 2019, 8 (7), 727. https://doi.org/10.3390/cells8070727.

(17) Ufer, F.; Vargas, P.; Engler, J. B.; Tintelnot, J.; Schattling, B.; Winkler, H.; Bauer, S.; Kursawe, N.; Willing, A.; Keminer, O.; Ohana, O.; Salinas-Riester, G.; Pless, O.; Kuhl, D.; Friese, M. A. Arc/Arg3.1 Governs Inflammatory Dendritic Cell Migration from the Skin and Thereby Controls T Cell Activation. Sci Immunol 2016, 1 (3), eaaf8665. https://doi.org/10.1126/sciimmunol.aaf8665.

(18) Rodríguez, J. J.; Davies, H. A.; Silva, A. T.; De Souza, I. E. J.; Peddie, C. J.; Colyer, F. M.; Lancashire, C. L.; Fine, A.; Errington, M. L.; Bliss, T. V. P.; Stewart, M. G. Long-Term Potentiation in the Rat Dentate Gyrus Is Associated with Enhanced Arc/Arg3.1 Protein Expression in Spines, Dendrites and Glia. European Journal of Neuroscience 2005, 21 (9), 2384–2396. https://doi.org/10.1111/j.1460-9568.2005.04068.x.

(19) Kremerskothen, J.; Wendholt, D.; Teber, I.; Barnekow, A. Insulin-Induced Expression of the Activity-Regulated Cytoskeleton-Associated Gene (ARC) in Human Neuroblastoma Cells Requires P21ras, Mitogen-Activated Protein Kinase/Extracellular Regulated Kinase and Src Tyrosine Kinases but Is Protein Kinase C-Independent. Neuroscience Letters 2002, 321 (3), 153–156. https://doi.org/10.1016/S0304-3940(01)02532-0.

(20) Korb, E.; Wilkinson, C. L.; Delgado, R. N.; Lovero, K. L.; Finkbeiner, S. Arc in the Nucleus Regulates PML-Dependent GluA1 Transcription and Homeostatic Plasticity. Nature neuroscience 2013, 16 (7), 874–883.

(21) Newpher, T. M.; Harris, S.; Pringle, J.; Hamilton, C.; Soderling, S. Regulation of Spine Structural Plasticity by Arc/Arg3.1. Semin. Cell Dev. Biol. 2018, 77, 25–32. https://doi.org/10.1016/j.semcdb.2017.09.022.

(22) Volkova, M. V.; Boyarintsev, V. V.; Trofimenko, A. V.; Biryukov, S. A.; Gorina, E. V.; Filkov, G. I.; Durymanov, M. O. Adaptation of Bio-Layer Interferometry for Quantitative Assessment of the Vascular Endothelial Growth Factor Content in Cell-Conditioned Culture Medium. Biophysics 2020, 65 (6), 935–941.

(23) Wang, Y.; Dzakah, E. E.; Kang, Y.; Cai, Y.; Wu, P.; Cui, Y.; Huang, Y.; He, X. Development of Anti-Müllerian Hormone Immunoassay Based on Biolayer Interferometry Technology. Anal Bioanal Chem 2019, 411 (21), 5499–5507. https://doi.org/10.1007/s00216-019-01928-6.

(24) Müller-Esparza, H.; Osorio-Valeriano, M.; Steube, N.; Thanbichler, M.; Randau, L. Bio-Layer Interferometry Analysis of the Target Binding Activity of CRISPR-Cas Effector Complexes. Front. Mol. Biosci. 2020, 7. https://doi.org/10.3389/fmolb.2020.00098.

(25) Vlassov, A. V.; Magdaleno, S.; Setterquist, R.; Conrad, R. Exosomes: Current Knowledge of Their Composition, Biological Functions, and Diagnostic and Therapeutic Potentials. Biochimica et Biophysica Acta (BBA)-General Subjects 2012, 1820 (7), 940–948.

(26) Dar, G. H.; Mendes, C. C.; Kuan, W.-L.; Speciale, A. A.; Conceição, M.; Görgens, A.; Uliyakina, I.; Lobo, M. J.; Lim, W. F.; EL Andaloussi, S.; Mäger, I.; Roberts, T. C.; Barker, R. A.; Goberdhan, D. C. I.; Wilson, C.; Wood, M. J. A. GAPDH Controls Extracellular Vesicle Biogenesis and Enhances the Therapeutic Potential of EV Mediated SiRNA Delivery to the Brain. Nat Commun 2021, 12 (1), 6666. https://doi.org/10.1038/s41467-021-27056-3.

(27) Ohgaki, H. Epidemiology of Brain Tumors. In Cancer Epidemiology: Modifiable Factors; Verma, M., Ed.; Methods in Molecular Biology; Humana Press: Totowa, NJ, 2009; pp 323–342. https://doi.org/10.1007/978-1-60327-492-0_14.

(28) Alcantara Llaguno, S. R.; Parada, L. F. Cell of Origin of Glioma: Biological and Clinical Implications. Br J Cancer 2016, 115 (12), 1445–1450. https://doi.org/10.1038/bjc.2016.354.

(29) Venkatesh, H. S.; Johung, T. B.; Caretti, V.; Noll, A.; Tang, Y.; Nagaraja, S.; Gibson, E. M.; Mount, C. W.; Polepalli, J.; Mitra, S. S.; Woo, P. J.; Malenka, R. C.; Vogel, H.; Bredel, M.; Mallick, P.; Monje, M. Neuronal Activity Promotes Glioma Growth through Neuroligin-3 Secretion. Cell 2015, 161 (4), 803–816. https://doi.org/10.1016/j.cell.2015.04.012.

(30) Venkataramani, V.; Tanev, D. I.; Strahle, C.; Studier-Fischer, A.; Fankhauser, L.; Kessler, T.; Körber, C.; Kardorff, M.; Ratliff, M.; Xie, R.; Horstmann, H.; Messer, M.; Paik, S. P.; Knabbe, J.; Sahm, F.; Kurz, F. T.; Acikgöz, A. A.; Herrmannsdörfer, F.; Agarwal, A.; Bergles, D. E.; Chalmers, A.; Miletic, H.; Turcan, S.; Mawrin, C.; Hänggi, D.; Liu, H.-K.; Wick, W.; Winkler, F.; Kuner, T. Glutamatergic Synaptic Input to Glioma Cells Drives Brain Tumour Progression. Nature 2019, 573 (7775), 532–538. https://doi.org/10.1038/s41586-019-1564-x.

(31) Hausmann, D.; Hoffmann, D. C.; Venkataramani, V.; Jung, E.; Horschitz, S.; Tetzlaff, S. K.; Jabali, A.; Hai, L.; Kessler, T.; Azoŕin, D. D.; Weil, S.; Kourtesakis, A.; Sievers, P.; Habel, A.; Breckwoldt, M. O.; Karreman, M. A.; Ratliff, M.; Messmer, J. M.; Yang, Y.; Reyhan, E.; Wendler, S.; Löb, C.; Mayer, C.; Figarella, K.; Osswald, M.; Solecki, G.; Sahm, F.; Garaschuk, O.; Kuner, T.; Koch, P.; Schlesner, M.; Wick, W.; Winkler, F. Autonomous Rhythmic Activity in Glioma Networks Drives Brain Tumour Growth. Nature 2023, 613 (7942), 179–186. https://doi.org/10.1038/s41586-022-05520-4.

(32) Perlman, M.; Resh, M. D. Identification of an Intracellular Trafficking and Assembly Pathway for HIV-1 Gag. Traffic 2006, 7 (6), 731–745. https://doi.org/10.1111/j.1398-9219.2006.00428.x.

(33) Sfakianos, J. N.; LaCasse, R. A.; Hunter, E. The M-PMV Cytoplasmic Targeting-Retention Signal Directs Nascent Gag Polypeptides to a Pericentriolar Region of the Cell. Traffic 2003, 4 (10), 660–670. https://doi.org/10.1034/j.1600-0854.2003.00125.x.

(34) Comas-Garcia, M.; Datta, S. A.; Baker, L.; Varma, R.; Gudla, P. R.; Rein, A. Dissection of Specific Binding of HIV-1 Gag to the “packaging Signal” in Viral RNA. eLife 2017, 6, e27055. https://doi.org/10.7554/eLife.27055.

(35) Royle, S. J.; Lagnado, L. Clathrin-Mediated Endocytosis at the Synaptic Terminal: Bridging the Gap Between Physiology and Molecules. Traffic 2010, 11 (12), 1489–1497. https://doi.org/10.1111/j.1600-0854.2010.01104.x.

(36) Preta, G.; Cronin, J. G.; Sheldon, I. M. Dynasore - Not Just a Dynamin Inhibitor. Cell Communication and Signaling 2015, 13 (1), 24. https://doi.org/10.1186/s12964-015-0102-1.

(37) Matejka, N.; Reindl, J. Influence of α-Particle Radiation on Intercellular Communication Networks of Tunneling Nanotubes in U87 Glioblastoma Cells. Frontiers in Oncology 2020, 10, 1691.

(38) Bramham, C. R.; Alme, M. N.; Bittins, M.; Kuipers, S. D.; Nair, R. R.; Pai, B.; Panja, D.; Schubert, M.; Soule, J.; Tiron, A.; Wibrand, K. The Arc of Synaptic Memory. Exp Brain Res 2010, 200 (2), 125–140. https://doi.org/10.1007/s00221-009-1959-2.

(39) Mikuni, T.; Uesaka, N.; Okuno, H.; Hirai, H.; Deisseroth, K.; Bito, H.; Kano, M. Arc/Arg3. 1 Is a Postsynaptic Mediator of Activity-Dependent Synapse Elimination in the Developing Cerebellum. Neuron 2013, 78 (6), 1024–1035.

(40) Ma, K.; Ding, X.; Song, Q.; Han, Z.; Yao, H.; Ding, J.; Hu, G. Lactate Enhances Arc/Arg3.1 Expression through Hydroxycarboxylic Acid Receptor 1-β-Arrestin2 Pathway in Astrocytes. Neuropharmacology 2020, 171, 108084. https://doi.org/10.1016/j.neuropharm.2020.108084.

(41) Caruso, J. P.; Koch, B. J.; Benson, P. D.; Varughese, E.; Monterey, M. D.; Lee, A. E.; Dave, A. M.; Kiousis, S.; Sloan, A. E.; Mathupala, S. P. PH, Lactate, and Hypoxia: Reciprocity in Regulating High-Affinity Monocarboxylate Transporter Expression in Glioblastoma. Neoplasia 2017, 19 (2), 121–134. https://doi.org/10.1016/j.neo.2016.12.011.

(42) Talacchi, A.; Santini, B.; Savazzi, S.; Gerosa, M. Cognitive Effects of Tumour and Surgical Treatment in Glioma Patients. J Neurooncol 2011, 103 (3), 541–549. https://doi.org/10.1007/s11060-010-0417-0

