## Supplementary material for "Retrovirus-Like Gag Protein Arc/Arg3.1 is Involved in Extracellular-Vesicle-Mediated mRNA Transfer between Glioma Cells": subblementary

**SUPPLEMENTARY METHODS**

**Gibson assembly**

The design of the primers was based on “classic” criteria such as avoiding regions of secondary structure and self-annealing, aiming at the GC content to be between 40 and 60 % with three C or G at the 3′ of a primer ending to promote binding. The PCR mixture contained 1× PCR buffer, 0.5 μM of each primer, 0.2 mM dNTPs mix and 0.02 U μL^−1^ of Q5® High-Fidelity DNA polymerase (New England BioLabs). The insert fragment encoding Arc-tGFP was amplified from pCMV6-AC-GFP plasmid containing Human Arc ORF Clone, using primers (Arc+GFP): Forward 5′-TCGCCACCTTTTGTAATACGACTCACTATAG -3′; Reverse 5′-GCGGCCGCTTAAACTCTTTCTTCACCG-3′. The fragment encoding the vector was amplified from eGFP Reporter Lentiviral plasmid pHAGE-CMV-eGFP using primers (lenti-plasmid): Forward 5′-AGTTTAAGCGGCCGCCATGGAATAAGG-3′; Reverse 5′-ATTACAAAAGGTGGCGACCGCTAGCGG -3′.

PCR reactions, containing 5–10 ng of the template plasmid, were carried out under the following conditions: the initial denaturation of 30 s at 98°C, followed by 35 cycles of 10s at 98°C, 30 s at 60°C, 40 s per kb at 72°C, followed by a final elongation of 2 min at 72°C. All amplified fragments were digested by the DpnI restriction enzyme (1× buffer, 0.5 U μL^-1^ of DpnI). Once DpnI recognizes 5′-GmeATC-3′ sites and digests the methylated DNA, it was used to degrade the plasmid template following the PCR reactions. Then, the amplified fragments were purified using the Cleanup Standard DNA extraction kit (Evrogen, Russia) according to the manufacturer’s instructions.

The two clean-up PCR fragments (insert and vector) were assembled according to the one-step isothermal DNA assembly method described by Gibson in 2009.^17^ Briefly, 0.05 pmol of DNA vector and 0.1 pmol of DNA insert were pooled in 3 μL, and 3 μL of Gibson assembly master mixture according to Gibson’s protocol (New England BioLabs) were added to the DNAs. The mixture was incubated at 50°C for 30 min in a thermocycler. To validate DNA assembly, 1 % agarose gel electrophoresis was performed in 1× TBE buffer and 5 μg mL^-1^ ethidium bromide for DNA visualization. Two microliters of Gibson assembly reaction were used to transform XL10-Gold ultracompetent *E.* *coli* cells according to the manufacturer’s recommendations (New Englands Biolabs). Bacterial plasmid DNA extraction was performed using Plasmid Miniprep kit (Evrogen, Russia) according to the manufacturer’s instructions. This new lentiviral construct containing Arc-GFP was called “plenti-Arc-GFP”.

***In vitro* production and biotinylation of target mRNAs**

Full-length mRNAs encoding human Arc protein and mCherry were obtained using the T7 RNA transcription kit (Biolabmix, Russia) in presence of DNA plasmids containing T7 promoter sequence. According to the supplier’s instruction, *in vitro* transcription reaction (50 µL) was performed at 37°C for 16 h. The reaction mixture contained 1 μg template DNA, 1 U µL^−1^ T7 polymerase, 1 mM NTP mixture, 1 μL of 25× DTT, 10 μL of 5× T7 RNA polymerase reaction buffer, and RNase-free water (up to 50 μL). Both RNAs were internally biotin-labelled by adding 0.5 mM of Biotin-11-UTP during the transcription reaction, at a molar ratio of modified NTP to standard NTP 1:2. The resulting reaction products were treated with 1 μL DNase I (RNase-free) for 20 min at 37°C. The transcript was purified using CleanRNA Standard (Evrogen, Moscow, Russia). The purified RNA was used immediately for the downstream reaction or stored at −80°C.

**EV fluorescent labeling**

The resuspended EVs were stained with PKH26 using PKH26 Red Fluorescent Cell Linker Kit for General Cell Membrane Labeling (Sigma Aldrich, USA). Labeling of U87-derived EVs with PKH26 was performed according to the manufacturer's protocol. Briefly, the EVs in 150 μL of PBS were mixed with 150 μL of 8 μM PKH26 solution (final PKH26 concentration is 4 μM) and incubated for 5 min at room temperature. For the control sample, particle-free Dulbecco's phosphate-buffered saline (DPBS; Sigma-Aldrich) was used as the input instead of the EV suspension. Then, the reaction was stopped by addition of 300 μL PBS containing 5 % BSA for 1 min to bind the excess. Then the EVs were diluted with DMEM and pelleted by ultracentrifugation at 100,000 × g for 70 min at 4 °C, washed with PBS and centrifuged again using the same parameters. The pellet was gently resuspended in 50 μL of DPBS. Incorporation of EVs into U87 cells was visualized by AxioVert.A1 microscope (Zeiss, Oberkochen, Germany) equipped with a ×20/0.6 objective lens after incubation with PKH26-labeled EVs for 6 h at 37^o^C.

**Generation of Arc knockout U87 cells**

To induce efficient knockout of the human Arc gene, different Arc-targeting sgRNAs were designed. Two free webtools to design sgRNAs were used (https://design.synthego.com and https://chopchop.cbu). Several factors have been considered when selecting a sgRNA from the databases. First, a 20mer sgRNA sequence with an intermediate GC content from 45 % to 65 % usually shows the highest efficiency. Second, the sgRNA should be located within the first 10–30 % of the first coding exon, rather than the last exon, to generate a shorter, loss-of-function truncated protein as early as possible. On this basis, three sgRNA sequences targeting the first exon were chosen to order annealing oligos. The following oligos have been used to introduce sgRNAs targeting Arc: (sgRNA1) upper: 5′-CACCCACCTGCGCACAGATGGAGC-3′; lower: 5′-AAACGCTCCATCTGTGCGCAGGTG-3′; (sgRNA2) upper: 5′-CACCGCCCGCCACACCGTTTCCGTGGG-3′; lower: 5′-AAACCCCACGGAAACGGTGTGGCGGGC-3′; (sgRNA3) upper: 5′-CACCCAGCGCTGCGAGTCGCTCGTGGG-3′; and lower: 5′-AAACCCCACGAGCGACTCGCAGCGCTG-3′. To generate the short version of pLenti-CRISPR/Cas9-expressing cassette, the plasmid lentiCRISPRv2 was treated with BsmBI restriction enzyme, separated by a short linker sequence, and cloned with a pair of annealed oligonucleotides. Colony PCR allowed for easy screening of sgRNA-positive clones. CRISPR clones were co-transfected with packaging vectors psPAX2 and pMD.2G in equimolar ratios into HEK293 cells to produce lentivirus. Production of lentivirus was performed following the author’s protocol. Lentiviral particles were concentrated by ultracentrifugation at 55,000 × g for 3 h at 4°C. The pellet was resuspended in DMEM, aliquoted and frozen at –80°C. The CRISPR/Cas9 lentivirus aliquots were used to transduce U87 cell line in 24-well plates, with different dilutions of the virus in each well (including no virus control). After puromycin selection (1 µg mL^-1^) for 4-7 days, survived infected cells have been used for further experiments. The quality of Arc KO U87 cells was examined by detection of Arc mRNA and Arc protein by qRT-PCR and WB, respectively.

**Immunocytochemistry (ICC)**

Glioma cells grown on glass coverslips for 24 h were washed 3 times with PBS, fixed for 20 min in 4 % paraformaldehyde, and permeabilized with 0.1 % Triton X-100/PBS for 5 min. Unspecific binding was blocked for 30 min with 1 % BSA in PBST. The cells were incubated with polyclonal rabbit anti-Arc antibodies (TA349500, OriGene) in 1 % BSA/PBST (dilution 1:500) for 1 h at RT, followed by 1 h incubation at RT with secondary antibodies (Abcam, ab150077) in 1 % BSA/PBST and 1 µg mL^-1^ DAPI. The actin cytoskeleton was stained using Phalloidin-iFluor 647 (Abcam, ab176759) for 1 h at RT (1:1000 dilution in 1 % BSA/PBS). The sample visualization was performed using EVOS M5000 microscope (Thermo Fisher Scientific, Carlsbad, CA).

**Cell viability assay**

The viability of U87 cells treated with 10 different concentrations (in a range of 10-200 µM) of endocytosis inhibitors was assessed by MTT assay. For all experiments, U87 cells were seeded to a 96-well plate at a density of 3,000 cells per well and incubated overnight before the treatment. The cells were treated with endocytosis inhibitors including nystatin (NYS) (Biosyntez, Penza, Russia), chlorpromazine (CPZ) (Novosibkhimpharm, Novosibirsk, Russia), and cytochalazine D (CytD) (L0418, Santa Cruz Biotechnology), and incubated at 37°C for 6 h. At the end of the incubation, the cells were washed three times with PBS and incubated with fresh supplemented media for an additional 24 h. MTT reagent (3-(4,5-dimethylthiazol-2-yl)-2,5-diphenyltetrazolium bromide) (ACROS Organics, Geel, Belgium) was added to the growth medium at a final concentration of 0.5 mg mL^-1^, followed by incubation at 37°C for 4 h. Supernatants were removed, and the cells were washed with PBS. The formazan crystals were dissolved in 200 μL dimethyl sulfoxide (DMSO, Sigma) for 10 min. Optical density was measured using CLARIOstar Plus spectrophotometer at 560 nm.

**SUPPLEMENTARY TABLES AND FIGURES**

**Table S1.** Parameters of Arc protein binding to mRNA*

|  | k_on_ (M^-1^s^-1^) ×10^5^ | k_off_ (s^-1^) ×10^-2^ | K_D_ (nM) | R^2^ |
| --- | --- | --- | --- | --- |
| Arc protein/*Arc* mRNA | 4.6 ± 0.6 | 3.6 ± 0.8 | 78 ± 11 | 0.95 |
| Arc protein/*mCherry* mRNA | 3.2 ± 0.3 | 3.5 ± 0.7 | 110 ± 19 | 0.95 |

* Data are shown as means ± SD.

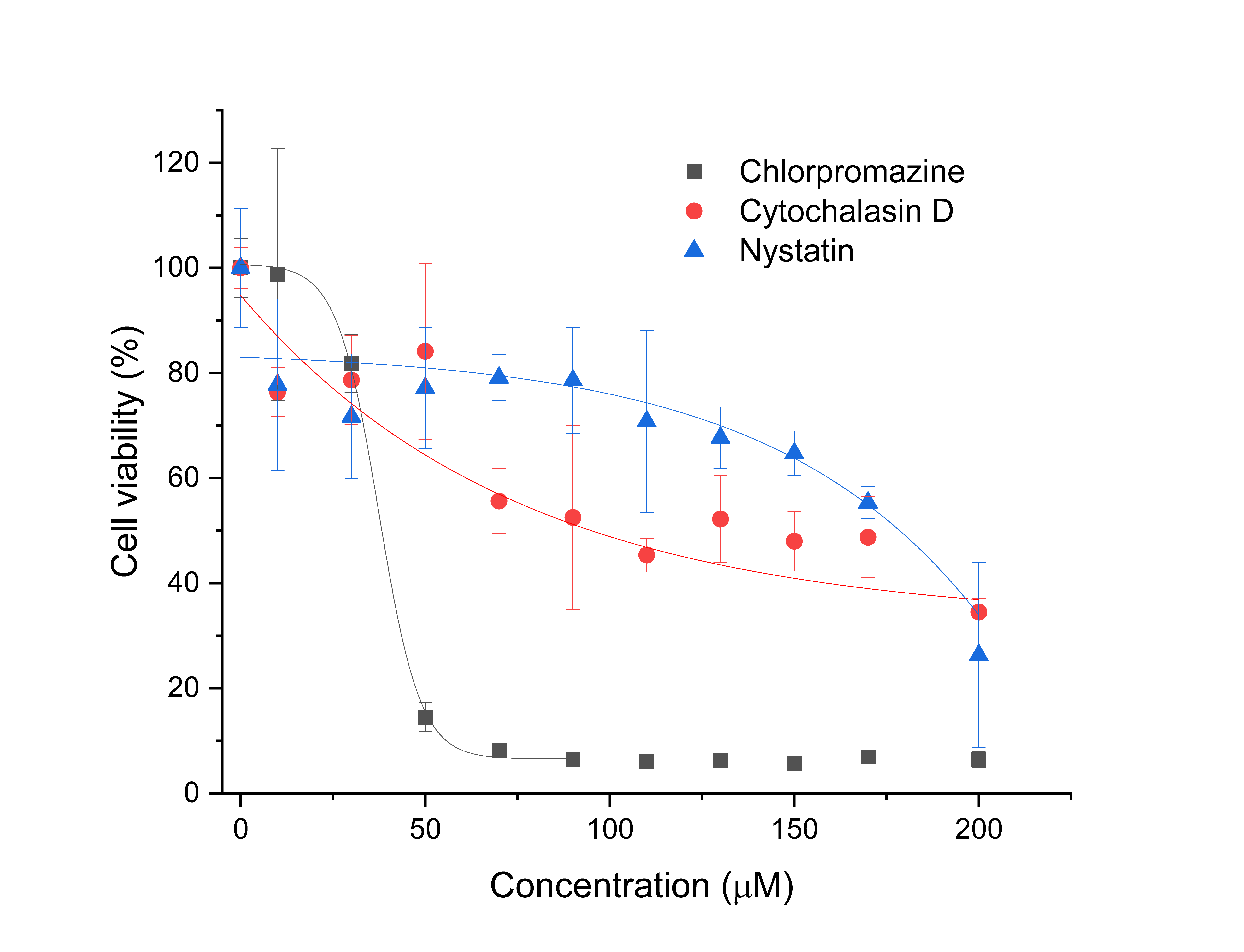

**Figure S1.** Analysis of U87 cell viability after treatment with pharmacological inhibitors of endocytic routes including chlorpromazine, nystatin, and cytochalasin D. The cells were incubated with inhibitors for 6 h, followed by 24-h incubation in growth medium. The data are given as means±SD.
